# A novel framework for single-cell Hi-C clustering based on graph-convolution-based imputation and two-phase-based feature extraction

**DOI:** 10.1101/2021.04.30.442215

**Authors:** Caiwei Zhen, Yuxian Wang, Lu Han, Jingyi Li, Jinghao Peng, Tao Wang, Jianye Hao, Xuequn Shang, Zhongyu Wei, Jiajie Peng

## Abstract

The three-dimensional genome structure plays a key role in cellular function and gene regulation. Singlecell Hi-C technology can capture genome structure information at the cell level, which provides the opportunity to study how genome structure varies among different cell types. However, few methods are well designed for single-cell Hi-C clustering, because of high sparsity, noise and heterogeneity of single-cell Hi-C data. In this manuscript, we propose a novel framework, named ScHiC-Rep, for singlecell Hi-C data representation and clustering. ScHiC-Rep mainly contains two parts: data imputation and feature extraction. In the imputation part, a novel imputation workflow is proposed, including graph convolution-based, random walk with restart-based and genomic neighbor-based imputation. In the feature extraction part, a two-phase feature extraction method is proposed, including linear phase for chromosome level and non-linear phase for cell level feature extraction. The evaluation results show that the proposed framework outperforms existing state-of-the-art approaches on both human and mouse datasets.

## Introduction

In recent years, there has been a rapid increase in the development of single-cell techniques^1^, including single-cell/nucleus RNA sequencing (RNA-seq)^2^, single-cell chromatin accessibility^3, 4^ and single-cell methylomes-based neuronal subtypes^5^. These powerful techniques provide opportunities to study the unique patterns of molecular characteristics that can distinguish different cell types. Based on various molecular characteristics, such as transcriptome^6, 7^, methylome^8^ and open chromatin^9–11^, different computational methods have been developed for cell clustering. Comparing with the bulk sequencing analysis, single-cell-based analysis introduces an opportunity to explore the biological mechanism at the cell level. However, unbiased and effective single-cell clustering algorithms based on the three-dimensional chromosome structure are still limited.

Three-dimensional genome structure has an important impact on gene regulation and cellular function. A large number of studies have shown that the spatial structure of chromosomes is closely related to most of the biological processes in cells, including genome regulation, stable expression of genome, DNA replication and chromosome ectopic^12–16^. Recently, chromosome conformation capture (3C) technology^17^ develops rapidly. 3C combines with Hi-C^18^ can capture genome-wide chromosome interaction maps^19^. Hi-C has the power of mapping the dynamic conformations of whole genomes^18^, which helps to analyze the hierarchical structure of chromosomes and reconstruct the three-dimensional structure^20^. Single-cell Hi-C^21^ technology is developed on the basis of Hi-C technology. Hi-C uses the averaged genome-wide interaction maps produced from bulk cells to evaluate chromosome folding and potential interactions^22^, while single-cell Hi-C can capture the genome structure information at cell level^21^. Single-cell Hi-C data enable researchers to reveal genome conformation patterns in different cell types and understand cell type heterogeneity^23^. However, because of intrinsic variability, data sparsity and coverage heterogeneity of single-cell Hi-C data^24^, it is still challenging to group cells based on single-cell Hi-C data.

Yardimci et al.^23^ have discussed whether the previously developed methods for analyzing bulk Hi-C data can be applied to single-cell Hi-C data. Several methods originally developed for bulk Hi-C data analysis have been applied for single-cell Hi-C data analysis^25^, such as HiCRep^26^, GenomeDISCO^27^, HiC-spector^28^ and QuASAR^29^. HiCRep takes the unique spatial features for the Hi-C data into account to evaluate the reproducibility of the data^26^. Firstly, HiCRep smooths the Hi-C map to decrease the effect of noise and biases. Secondly, HiCRep stratifies Hi-C map on the basis of the genomic distance to minimize the impact of distance dependence. Then, HiCRep evaluates the reproducibility of Hi-C maps from different cell types by proposing a stratum adjusted correlation coefficient (SCC). GenomeDISCO can be used to calculate the similarity of the two contact maps generated from the 3C experiment^27^. The key idea of this method is using random walk to impute the contact map and then calculating the concordance score between the imputed contact maps. Biological replicates and samples generated from different cell types can be separated accurately by GenomeDISCO. Based on spectral decomposition, a novel reproducibility metric to quantify the similarity between contact maps has proposed by HiC-Spector^28^. Contact maps from different biological replicates, pseudo-replicates and different cell types can be separated successfully by the metric.

The differentiation of different cell types is vital for the study of immunology, oncology, genetics and cell subsets, which can improve our understanding of the role of genomic structure in gene regulation in specific cell types. However, there are few methods designed for single-cell Hi-C clustering. Combining with multidimensional scaling (MDS) method, HiCRep, GenomeDISCO and HiC-Spector can be used to analyze single-cell Hi-C data. Yadimic et al.^23^ have shown that HiCRep performs significantly better than GenomeDISCO and HiC-Spector when embedding single-cell Hi-C data into two dimensions. The low dimensional representation of the cell similarity based on MDS is usually continuous. These MDS-based methods cannot perform well on cell clustering based on single-cell Hi-C data. Zhou et al. proposed a computational framework, named scHiCluster^24^. Firstly, it uses linear convolution and random walk with restart to impute the contact matrix. Then it selects the top 20% interactions and uses Principal Component Analysis (PCA) for dimension reduction. Finally, the first two Principal Components (PCs) are plotted to visualize the cells and the first ten PCs are used for K-means++ clustering. The results show that the performance of scHiCluster is better than HiCRep combining with MDS and GenomeDISCO combining with MDS.

Although scHiCluster can improve the performance of sing-cell Hi-C clustering, the two-dimensional convolution is designed for image data originally. By applying to the Hi-C contact matrix, it can only capture the local information between genomic bins in near genomic distance. However, it has been shown that long-range chromatin interactions are also important in gene regulation^30^. Furthermore, since only PCA is used for dimension reduction and feature extraction, scHiCluster cannot capture non-linear dependency in the input data.

In the past few years, graph representation learning-based method has achieved great success in bioinformatics field^31, 32^. The essence of performing graph convolution on the networks is to achieve feature aggregation between related nodes. The Hi-C contact probability map (contact distance profile) can be considered as a network. The node is fixed-width genomic loci (typically using bin to describe nodes). The edge is the relation between loci *i* and loci *j* (typically using the number of observed reads as the weight of edges). By considering the Hi-C contact map as a network, we use graph convolution-based method to impute contact matrices for better capturing the topological structures. The graph convolution-based imputation can capture not only local interactions but also long-range chromatin interactions. To extract the non-linear features of single-cell Hi-C data, we develop a two-phase feature extraction method for single-cell Hi-C data by combining one linear and one non-linear feature-extraction process. The proposed feature extraction method mainly contained two components: linear phase based on PCA and non-linear phase based on AE.

In this manuscript, we present a novel graph convolution-based framework, named ScHiC-Rep, to cluster single cells based on Hi-C contact matrices. Here are the major contributions:

- A novel imputation workflow for single-cell Hi-C contact matrix is proposed, including graph convolution-based, random walk with restart-based and genomic neighbor-based imputation.
- A two-phase feature extraction method is proposed for learning the feature representation of a cell based on imputed single-cell Hi-C contact matrix, including linear phase for chromosome level and non-linear phase for cell level feature extraction.
- The evaluation results show that the proposed framework outperforms existing state-of-the-art approaches.

## Results

### Overview of ScHiC-Rep

We propose a novel computational framework called ScHiC-Rep for single-cell clustering based on single-cell Hi-C data. The framework of ScHiC-Rep is shown in Figure 1. It mainly contains two parts. First, given a contact map of each chromosome, we apply several imputation steps to obtain an imputed contact matrix (Fig. 1a). Second, a two-phase feature extraction model is proposed to extract the feature representation of each cell based on the imputed contact matrix (Fig. 1b). The extracted feature of each cell can be used for further clustering and visualization. The sparsity of single-cell Hi-C data is usually high because of experimental limitations of material dropout^23, 24^.

**Figure 1.**
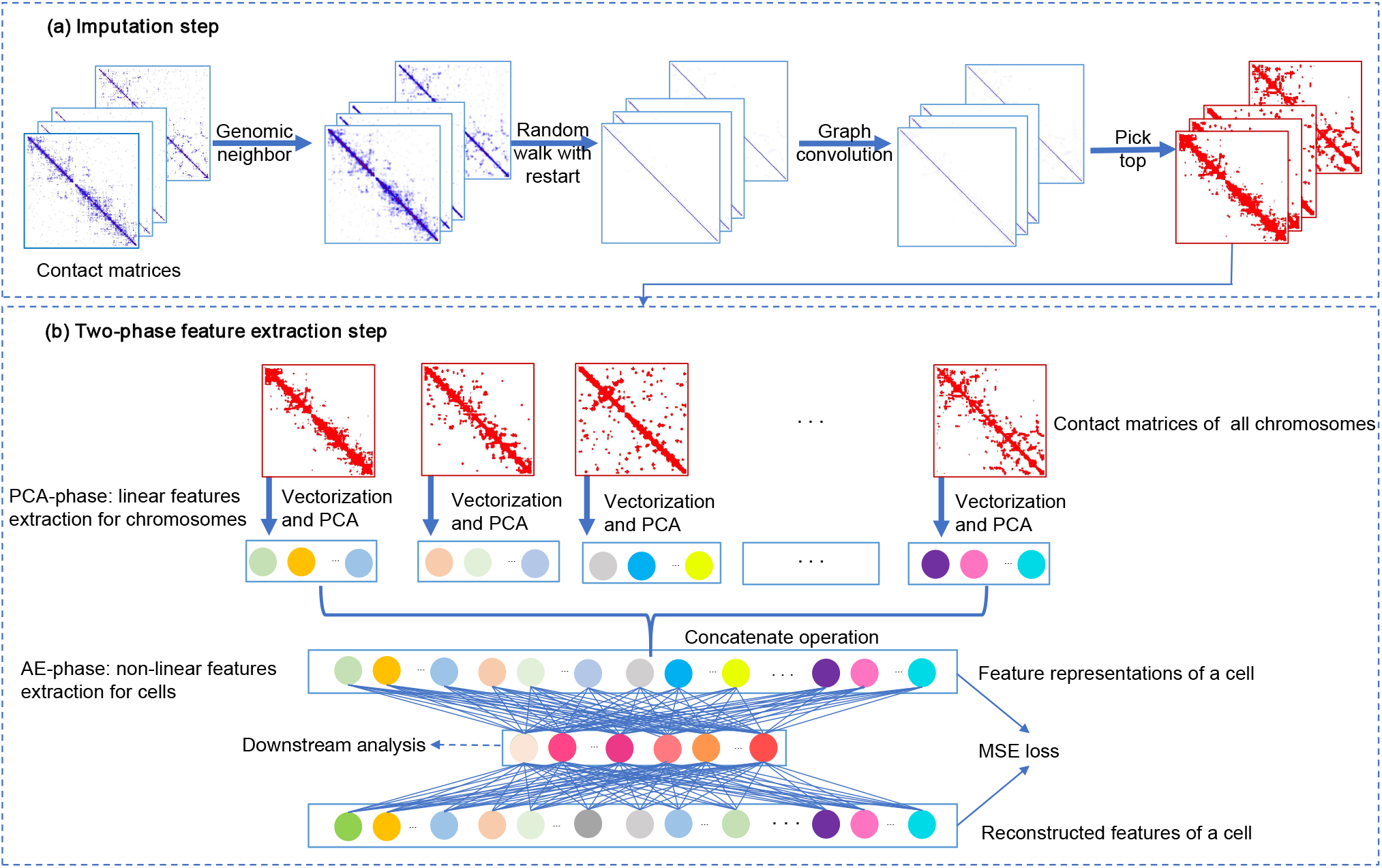
The workflow of ScHiC-Rep. The proposed method contains two main steps: (a). Given the contact matrices of all chromosomes as input, we first use genomic neighbor, random walk with restart and graph convolution-based method to impute the contact matrices. Then we pick top elements to generate binary matrices. (b). The feature extraction contains two-phase. In PCA-phase, given the binary matrices as input, we first generate the features of every chromosome and use PCA to generate the embedding of chromosomes. In AE-phase, we use concatenate operation to generate the feature representations of cells, then we use an AE to generate the low-dimensional representations of cells. Finally, the low-dimensional representations can be used for further downstream analysis, such as clustering and visualization.

The high sparsity usually leads to inaccurate downstream data analysis. To address this problem, we develop a computational method for single-cell Hi-C data imputation based on graph convolution method. The proposed workflow includes three steps: (1) genomic neighbor-based imputation; (2) random walk with restart-based imputation; (3) graph convolution-based imputation.

The two-phase feature extraction method mainly includes two components: linear feature extraction at chromosome level and non-linear feature extraction at cell level. On one hand, the feature dimension of contact matrix for chromosomes is extremely high. Furthermore, the features usually contain noisy and redundant data. Therefore, at the linear phase, a PCA^33^-based method is proposed to reduce the dimensionality of the feature space at chromosome level. However, PCA can only learn the linear features. In order to enhance the feature representation, we adopt autoencoder (AE) to learn the non-linear feature combinations at cell level.

On other hand, as a result of the features of chromosomes are extremely sparse and high dimensional, it is difficult to train an AE model by taking the contact matrix as input. The proposed model can obtain the dense features from high dimensional and sparse input features, and perform a deep neural network-based model to learn the non-linear and unseen feature combinations of a cell based on the dense chromosome level features.

### Performance evaluation on Flyamer Dataset

Same as the scHiCluster, we tested the utility of our computational framework on the Flyamer dataset. The experiment results show that ScHiC-Rep achieves the highest performance among all methods according to ARI, NMI, HM, and FM on Flyamer dataset. The average ARI score across 50 runs of ScHiC-Rep is 0.8049, which is significantly higher than the scores of FastICA, NMF, PCA, SVD, scHiCluster and Decay (Figure 2 and Supplementary Document Table S4). It is noted that the average ARI of ScHiC-Rep is about 0.1 higher than the second best method Decay. Similarly, ScHiC-Rep achieves the highest NMI (0.7166), which is about 0.08 higher than the second-best method scHiCluster. The average HM of ScHiC-Rep (0.7520) is also significantly higher than the second-best method scHiCluster. It is also shown that ScHiC-Rep can achieve the highest FM (0.9058). In summary, this experiment shows that ScHiC-Rep can achieve significant improvement on Flyamer dataset comparing with the state-of-art methods.

**Figure 2.**
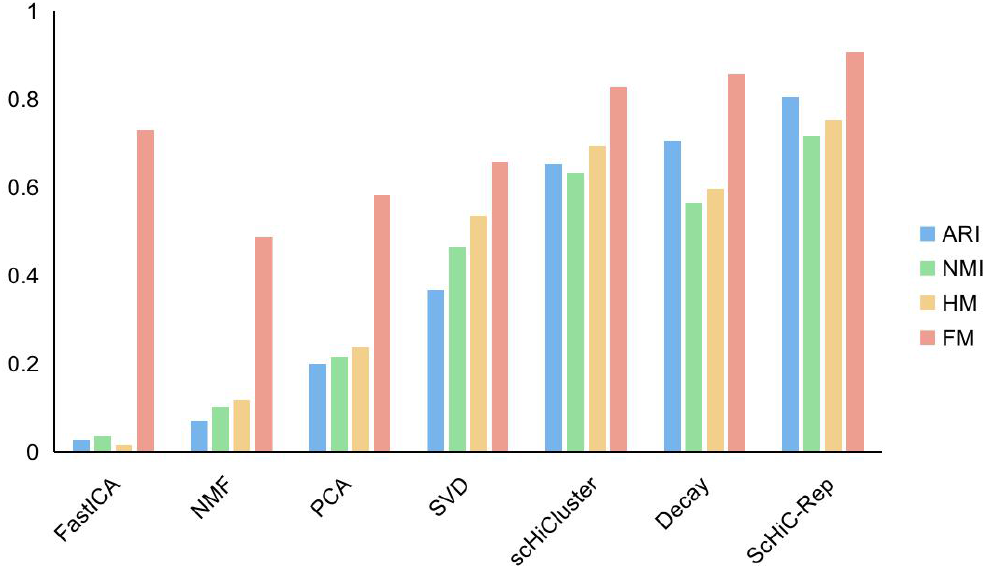
Quantitative evaluation of the ScHiC-Rep on Flyamer dataset. Compared with the seven methods, the ARI, NMI, HM and FM of ScHiC-Rep are both highest.

### Performance evaluation on Ramani Dataset

We also evaluate the proposed method on Ramani Dataset. The experiment results show that ScHiC-Rep achieves the best performance among all compared methods according to ARI, NMI, HM, and FM on Ramani dataset. The average score of ARI achieved by ScHiC-Rep across 10 runs is 0.8822, which is higher than the scores of FastICA, NMF, PCA, SVD, scHiCluster and Decay (Figure 3 and Supplementary Document Table S5). Comparing the NMI from the seven methods shows that ScHiC-Rep performs the best among them. The average NMI of ScHiC-Rep is about 0.015 higher than the second-best method scHiCluster. Comparing the HM with the seven methods shows that ScHiC-Rep performs the best. The average HM of ScHiC-Rep is about 0.08 higher than the second-best method scHiCluster. It is also shown that ScHiC-Rep can achieve the highest FM (0.9264). In summary, ScHiC-Rep performs better than some state-of-the-art methods on Ramani dataset.

**Figure 3.**
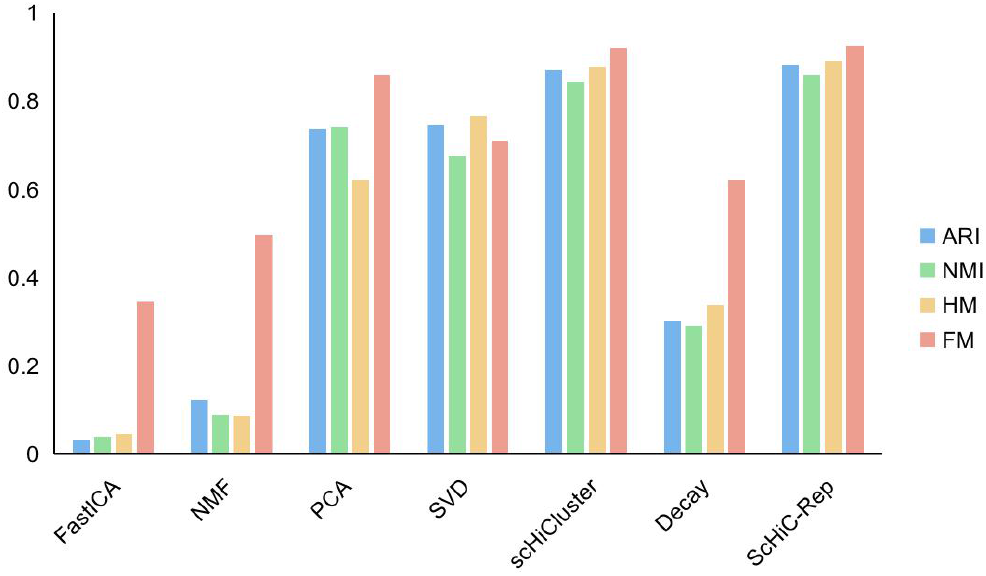
Quantitative evaluation of the ScHiC-Rep on Ramani dataset. Compared with the seven methods, the ARI, NMI, HM and FM of ScHiC-Rep are both highest.

### Effects of ScHiC-Rep components

In order to evaluate the effects of two components of ScHiC-Rep, we compare ScHiC-Rep with the other two alternative versions. To test the effect of graph convolution component in the proposed method, we test an alternative model named CR-PCAAE. In CR-PCAAE model, we removed the graph convolution-based step from the imputation workflow. To test the effect of the feature extraction step, we test an alternative model named GC-PCA. In GC-PCA model, the AE model (non-linear feature extraction phase) is replaced by PCA model. Table 1 shows that ScHiC-Rep performs better than CR-PCAAE and GC-PCA in terms of all metrics, indicating that the two components in ScHiC-Rep contribute to the performance and have been appropriately designed.

**Table 1.** The ARI, NMI, HM, and FM of GC-PCA and CR-PCAAE on Flyamer dataset and Ramani dataset. Bolded numbers are the best performance in each category.

| Dataset | Method | ARI | NMI | HM | FM |
| --- | --- | --- | --- | --- | --- |
| Flyamer dataset | GC-PCA | 0.7791 | 0.7250 | 0.7696 | 0.8936 |
|  | CR-PCAAE | 0.7964 | 0.7068 | 0.7413 | 0.9024 |
|  | ScHiC-Rep | **0.8715** | **0.8358** | **0.8700** | **0.9201** |
| Ramani dataset | GC-PCA | 0.8715 | 0.8358 | 0.8700 | 0.9201 |
|  | CR-PCAAE | 0.8803 | 0.8565 | 0.8893 | 0.9256 |
|  | ScHiC-Rep | **0.8822** | **0.8583** | **0.8913** | **0.9264** |

### Visualization of the learned embedding on two datasets

For the Flyamer dataset, as shown in Figure 4, ScHiC-Rep performs better than scHiCluster in three visualization methods. Comparing with scHiCluster, projection of embedding vector based on ScHiC-Rep into PC1 and PC2 results in closer Oocyte cells (Fig. 4a and Fig. 4d). Using tSNE to visualize the embedding vector (Fig. 4b and Fig. 4e), ScHiC-Rep makes both three types of cells more compact comparing with scHiCluster. Similar result is showin in Fig. 4c and Fig. 4f for UMAP-based visulization.The visualization and performance evaluation (Supplementary Documen Table S4) indicate the effectiveness of ScHiC-Rep, which has great potential to provide insights into the cell-to-cell variation of higher-order genome organization.

**Figure 4.**
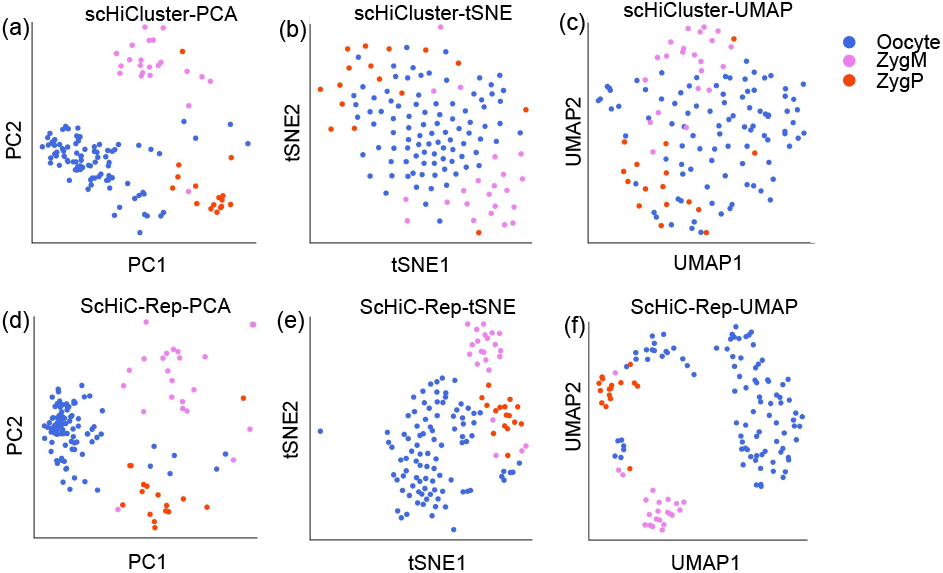
The visualization of the learned embedding based on scHiCluster on Flyamer dataset using PCA (a), tSNE (b) and UMAP (c). The visualization of the learned embedding based on ScHiC-Rep on the Flyamer dataset using PCA (d), tSNE (e) and UMAP (f).

For the Ramani dataset (Figure 5), ScHiC-Rep also performs better than scHiCluster in three visualization methods. Projection of embedding vectors based on ScHiC-Rep into PC1 and PC2 results in separation of K562 cells from HAP1 cells and GM12878 cells comparing with scHiCluster (Fig. 5a and Fig. 5d). In the task of using tSNE to visualize the embedding vector (Fig. 5b and Fig. 5e), ScHiC-Rep also separates K562 cells from HAP1 cells and GM12878 cells. Projection of embedding vectors of ScHiC-Rep into UMAP1 and UMAP2 results in more compact HeLa cells and K562 cells (Fig. 5c and Fig. 5f). HAP1, GM12878, and K562 cells are not clearly separated as cells in Ramani dataset. The reason might be that all these cells are blood-related cells^34^. Their 3D genome organizations are likely to be similar to each other. Quantitative results in Supplementary Document Table S5 are consistent with the visualization as ScHiC-Rep achieves higher ARI, NMF, HM, and FM score comparing with scHiCluster. We can find the visualization results for Decay and PCA methods on Flyamer dataset and Ramani dataset in Supplementary Document Figure S4-6.

**Figure 5.**
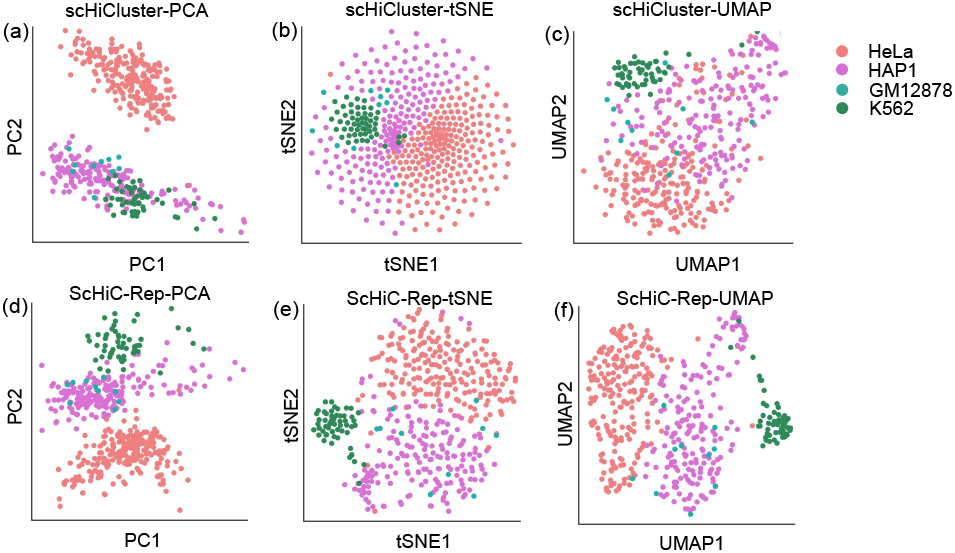
The visualization of the learned embedding based on scHiCluster on Ramani dataset using PCA (a), tSNE (b) and UMAP (c). The visualization of the learned embedding based on ScHiC-Rep on the Ramani dataset using PCA (d), tSNE (e) and UMAP (f).

## Conclusion

Recently, researchers have started to study cell-to-cell variation based on single-cell Hi-C data. In this manuscript, we propose a novel framework for single-cell clustering based on single-cell Hi-C data. The proposed framework mainly includes two parts. We first generate high-quality contact matrix based on graph convolution-based imputation, random walk with restart-based imputation and genomic neighbor-based imputation. Then, a two-phase feature extraction method is proposed to learn feature representation of a cell based on the imputed matrix.

After obtaining the feature representation of each cell, we apply K-means++ as the clustering algorithm for single-cell clustering. To evaluate the performance of ScHiC-Rep, we compare it with six methods. The evaluation results on both Flyamer dataset and Ramani dataset show that ScHiC-Rep performs better than existing methods, indicating that the proposed imputation method and two-phase feature extraction-based method is appropriately designed. In the future, we will develop a web server and include more datasets for convenient use of ScHiC-Rep.

## Materials and methods

### Single-cell Hi-C data imputation

#### Genomic neighbor-based imputation

The single-cell Hi-C data is often represented as a contact matrix. The contact matrix is a symmetry matrix which describes the interactions between genomic bins. We can represent the contact matrix as a network 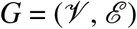, where 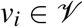 represents the node in the network, and 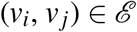 represents the edge in the network. In detail, where rows and columns represent the number of read-pairs that support the interaction between one chromosome and other chromosome fragments. The order of the rows and columns is usually based on the genomic positions.

In this step, aggregating the interaction information of the genomic neighbors for each contact is used to capture the local domain structure. Because of the dropout problem, the sparsity of contact matrix is usually high, so the 0 value in contact matrix may also indicate that there have interactions at that position. Consequently, genomic neighbor-based imputation method^24^ is proposed based on ‘guilty by association’. More specifically, given two bins *b*_1_ and *b*_2_, if *b*_1_ is close to *b*_2_, *b*_1_ may also be close to neighbors of *b*_2_. Based on this hypothesis, we can impute the missing values by capturing the local signals.

Specifically, a *m × m* kernel is used to scan a contact matrix 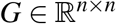, where *m* = 2*w* + 1 and *w* is the window size. The parameter selection of *w* is described in Supplementary Document Figure S1.

Given window size w, we apply a filter *F* of size *m* × *m* (where *m* =2*w* +1) to scan a contact matrix *G* of size *n* × *n*. Let 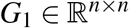 be the imputed contact matrix. The elements in the matrix *G*_1_ are imputed based on their neighbors as follows.

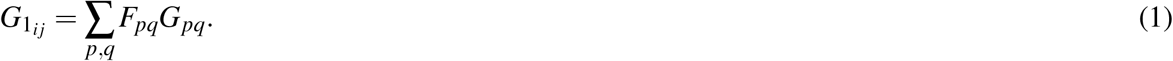

where *i* − *w* ≤ *p* ≤ *i* + *w*, *j* − *w* ≤ *p* ≤ *j* + *w*. The *F* is a all-one matrix. *w* is an adjustable threshold, which determines the number of neighbors considered. After this step, the input contact matrix is imputed based on the genomic neighbor information.

#### Random walk with restart-based imputation

After the genomic neighbor-based imputation, we can get a matrix *G*_1_. Taking *G*_1_ as input, we use a random walk with restart-based method^24^ to impute the contact matrix. Different from genomic neighbor-based imputation, the contact matrix is imputed based on the global domain structures at this step. RWR is used to calculate the probability of each pair of nodes from node *i* to node *j*, the final probability is calculated after *t*-th iteration. Therefore, the global signal of the network is taken into account.

First, the input contact matrix is normalized by row as follows:

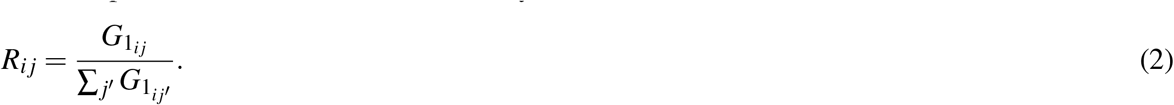

Then, given 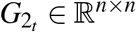 representing the matrix after the *t*-th iteration in RWR, *G*_2_*t*__ can be calculated recursively as follows:

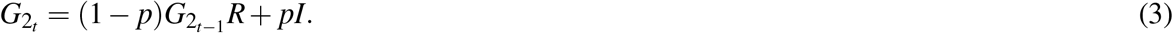

where *p* represents the restart probability, which can be used to balance the information between global domain structure and local domain structure. The parameter selection of *p* is described in Supplementary Document Figure S2. We initialize an identity matrix *G*_2_0__ =*I*, and iteratively perform RWR algorithm until ∥*G*_2_*t*__ − *G*_2_*t*−1__∥_2_ ≤ 10^−6^. Then, we can get an imputed matrix *G*_2_. *G*_2_*ij*__ represents the random walk probability from *v_i_* to *v_j_*, which represents the interaction between corresponding bin *i* to *j* in the contact matrix.

#### Graph convolution-based imputation

The contact matrix can be considered as a network describing the interaction between genomic bins. Graph convolution network (GCN) is a powerful technique for network analysis. GCN performs well for analyzing several types of networks, such as citation networks^35^, social networks^36^ and biological networks^37^.

Inspired by the success of GCN in network analysis, we propose a graph convolution-based method for contact matrix imputation. Then, the graph convolution-based imputation method is applied to aggregate the interaction information from one-hop neighbors. In traditional GCN model, two types of input data are needed. One is the feature of each node. The other is the network structure. By propagating the features of neighbor nodes, GCN can learn better feature representation of nodes.

In our work, for transferring node feature learning problem to adjacent matrix imputation problem, we use the *G*_2_ as the feature matrix and also the adjacent matrix network (network structure). We first calculate the normalized adjacency matrix *L* of *G*_2_ as follows:

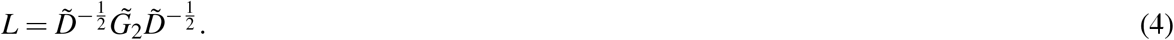

where 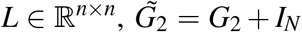. In this way, the adjacency matrix 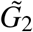 not only contains connections between nodes but also the information of each node itself. 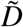 is the degree matrix of 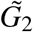. 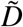 is defined as 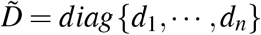, where each element on the diagonal is equal to the row-sum of the adjacency matrix, *d_i_* = ∑_*j*_ *G*_2_*ij*__.

Let the feature matrix be *X* = *G*_2_. The feature of a genomic bin *v_i_* is represented as *x_i_*, which is a feature vector. The initial value of *x_i_* is the *i*-th row of *X*. Then, the feature of *x_i_* is calculated as follows:

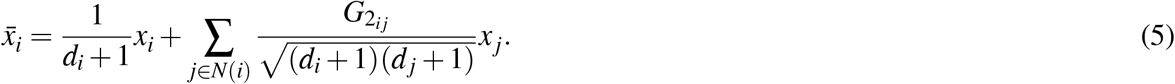

where *x_i_* represents the initial feature of *v_i_*, *x_j_* represents the initial feature of *v_j_*. *v_j_* is the neighbor of *v_i_*. *N*(*i*) represents the one-hop neighbors set of 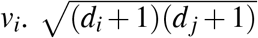 is the symmetric sqrt normalization constant. *d_i_* and *d_j_* represent the degree of *v_i_* and *v_j_* respectively. 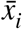 represents the feature of *v_i_* after aggregating the feature of its one-hop neighbors.

More intuitively, the above process can be simplified as follows:

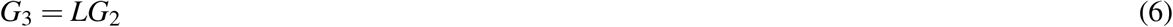

where 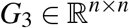, *L* is the normalized adjacency matrix of *G*_2_. *G*_3_ is the imputed contact matrix, which is the input of feature extraction step. Comparing the contact matrix before and after graph convolution-based imputation, the distributions of values in the contact matrix are significantly different (Supplementary Document Figure S7-10 and Supplementary Document Table S6-11).

### Two-phase feature extraction

In previous step, we obtain the imputed contact matrix for each chromosome of a cell. In this part, we propose a two-phase feature extraction method to learn the feature representation of a cell.

#### Feature extraction at chromosome level

Given an imputed contact matrix *G*_3_ of a chromosome, we keep the top *k* interactions to reduce the bias caused by uneven sequence coverage^24^. *k* is an adjustable threshold. Since the coverage of the contact matrix of different cell lines are different, the sparseness gap and the value gap between the matrices after imputation processing are also obvious. Therefore, after the imputation process, given a threshold *t*, we convert the imputed contact matrix *G*_3_ into a binary matrix *G_b_*. *G_b_* can be calculated as follows:

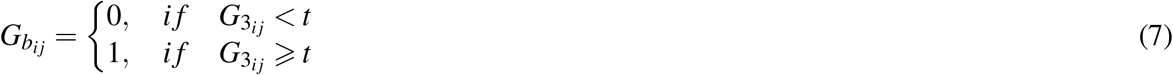

where *t* is an adjustable threshold, in this manuscript, *t* is set to be the 85-th percentile of values in *G_b_*. The parameter evaluation of *t* is described in Supplementary Document Figure S3. Then, a flatten operation is used to generate the chromosome feature vector 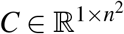, which can be described as:

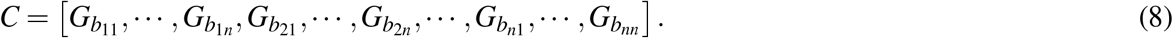

Taking 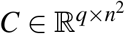 as input, we use PCA to reduce the dimension of the chromosome feature. *q* is the number of cells. A lower-dimension feature 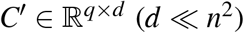 is generated. PCA is performed on each chromosome to get the lower-dimension feature representation. For each cell, we concatenate low-dimension features of all chromosomes to generate the feature of a cell. In detail, we use PCA to extract features in the same chromosome of all cells at the same time. The feature vector of a cell is represented as 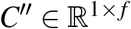 is, where *k* is the number of chromosomes, *f* = *d* ∗ *k*. Finally, we can get a feature matrix of all cells 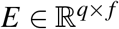.

In summary, PCA is used to project the chromosome contact matrix into a low-dimensional space and learn the linear feature combination. After PCA dimension reduction, all feature vectors of chromosomes are concatenated into a feature vector of a cell.

#### Autoencoder-based feature extraction at cell level

In this step, we use AE-based model^38^ to get the feature representation of each cell.

The neural network structure of AE model mainly contains two parts: data compression and data reconstruction.

In the data compression part, given a feature matrix of all cells 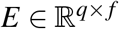, where *q* is the number of cells and *f* is the number of features. We use two hidden layers to compress the feature matrix. Then, a low-dimensional feature matrix *H* is generated. Given the feature matrix *H*, we use two layers neural network for reconstruction and generate a matrix 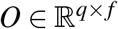. Mean Square Error (*MSE*) is used as loss function to optimize the matrix *H*. The loss function can be described as follows:

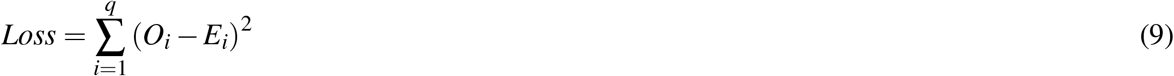

where *E* and *O* represent the original feature matrix and the reconstructed feature matrix respectively.

In the proposed model, we use the mean-square error (MSE)^39^ as the loss function. The ReLU activation function (on Flyamer dataset)^40^, Tanh activation function (on Ramani dataset)^41^ and Adam algorithm^42^ are used to optimize the model. The proposed model is trained by the backpropagation (BP) algorithm^43^. We use the grid search to find the best combination of parameters. The details can be found in Supplementary Document Table S2 and Table S3. Finally, given the low-dimensional feature matrix of all cells *H*, we use K-means++^44^ to cluster the cells.

### Experiment setup

We evaluate the proposed method on two datasets, namely, Ramani dataset^45^ and Flyamer dataset^46^, which are also used in previous studies^23, 24^. Given the single-cell Hi-C data, we first learn the feature representation of cells based on the proposed method and then use K-means++ as the clustering algorithm to group the cells. We use adjusted rand index (ARI)^47^, normalized mutual information (NMI)^48^, homogeneity (HM)^49^ score and fowlkes mallows (FM)^50^ score as the clustering result evaluation metrics.

We compare the performance of ScHiC-Rep with the other six methods, including FastICA^51^, Decay^52^, NMF^53^, SVD^54^, PCA^33^ and scHiCluster^24^ on the task of single-cell Hi-C clustering.

### Data description

Currently, single-cell Hi-C data is generally represented as two-dimensional contact matrix. The data generated by high-throughput sequencing technology such as Hi-C can be used to construct a contact matrix after read mapping and data filtering. The contact matrix generated by Hi-C experiment represents the interaction information between any genomic bins.

We evaluate our model on Ramani dataset^45^ and Flyamer dataset^46^. The Ramani dataset^45^ is downloaded from GSE84920, and the corresponding chromosome file can be downloaded from (http://genome.ucsc.edu/cgi-bin/hgTables). This dataset contains four human cell lines: HAP1, GM12878, K562, and HeLa. The Flyamer dataset^46^ is downloaded from GSE80006. Flyamer et al. generated files for the experimental cells with a resolution of 40-kbp and 200-kbp to save the contact matrix data information. We downloaded the 200-kbp data and merged the data into 1-Mbp data for single-cell Hi-C clustering experiments. Flyamer dataset contains three mouse cell lines: Oocyte, ZygM and ZygP. More details about the two datasets can be found in Supplementary Document Table S1. ScHic-Rep performs the same quality control on cells as scHiCluster^24^.

## Code availability

The source will be available after the manuscript is accepted.

## References

1. Tanay, A. & Regev, A. Scaling single-cell genomics from phenomenology to mechanism. Nature 541, 331–338 (2017).

2. Ramsköld, D. et al. Full-length mrna-seq from single-cell levels of rna and individual circulating tumor cells. Nat. biotechnology 30, 777–782 (2012).

3. Cusanovich, D. A. et al. Multiplex single-cell profiling of chromatin accessibility by combinatorial cellular indexing. Science 348, 910–914 (2015).

4. Buenrostro, J. D. et al. Single-cell chromatin accessibility reveals principles of regulatory variation. Nature 523, 486–490 (2015).

5. Luo, C. et al. Single-cell methylomes identify neuronal subtypes and regulatory elements in mammalian cortex. Science 357, 600–604 (2017).

6. Levine, J. H. et al. Data-driven phenotypic dissection of aml reveals progenitor-like cells that correlate with prognosis. Cell 162, 184–197 (2015).

7. Macosko, E. Z. et al. Highly parallel genome-wide expression profiling of individual cells using nanoliter droplets. Cell 161, 1202–1214 (2015).

8. Luo, C. et al. Robust single-cell dna methylome profiling with snmc-seq2. Nat. communications 9, 1–6 (2018).

9. Schep, A. N., Wu, B., Buenrostro, J. D. & Greenleaf, W. J. chromvar: inferring transcription-factor-associated accessibility from single-cell epigenomic data. Nat. methods 14, 975–978 (2017).

10. Cusanovich, D. A. et al. The cis-regulatory dynamics of embryonic development at single-cell resolution. Nature 555, 538–542 (2018).

11. Preissl, S. et al. Single-nucleus analysis of accessible chromatin in developing mouse forebrain reveals cell-type-specific transcriptional regulation. Nat. neuroscience 21, 432–439 (2018).

12. Misteli, T. Spatial positioning: A new dimension in genome function. Cell 119, 153–156 (2004).

13. Dekker, J. Gene regulation in the third dimension. Science 319, 1793–1794 (2008).

14. Miele, A. & Dekker, J. Long-range chromosomal interactions and gene regulation. Mol. biosystems 4, 1046–1057 (2008).

15. Fraser, P. & Bickmore, W. Nuclear organization of the genome and the potential for gene regulation. Nature 447, 413–417 (2007).

16. Alt, F. W., Zhang, Y., Meng, F.-L., Guo, C. & Schwer, B. Mechanisms of programmed dna lesions and genomic instability in the immune system. Cell 152, 417–429 (2013).

17. Dekker, J., Rippe, K., Dekker, M. & Kleckner, N. Capturing chromosome conformation. science 295, 1306–1311 (2002).

18. Lieberman-Aiden, E. et al. Comprehensive mapping of long-range interactions reveals folding principles of the human genome. science 326, 289–293 (2009).

19. Zhang, X.-Y. et al. Optimization and quality control of genome-wide hi-c library preparation. Yi Chuan= Hered. 39, 847–855 (2017).

20. Gao, J. et al. Developing bioimaging and quantitative methods to study 3d genome. Quant. Biol. 4, 129–147 (2016).

21. Nagano, T. et al. Single-cell hi-c reveals cell-to-cell variability in chromosome structure. Nature 502, 59–64 (2013).

22. Dekker, J. & Mirny, L. Chromosomes captured one by one. Nature 502, 45–46 (2013).

23. Liu, J., Lin, D., Yardimci, G. G. & Noble, W. S. Unsupervised embedding of single-cell hi-c data. Bioinformatics 34, i96–i104 (2018).

24. Zhou, J. et al. Robust single-cell hi-c clustering by convolution-and random-walk–based imputation. Proc. Natl. Acad. Sci. 116, 14011–14018 (2019).

25. Yardimci, G. G. et al. Measuring the reproducibility and quality of hi-c data. Genome biology 20, 1–19 (2019).

26. Yang, T. et al. Hicrep: assessing the reproducibility of hi-c data using a stratum-adjusted correlation coefficient. Genome research 27, 1939–1949 (2017).

27. Ursu, O. et al. Genomedisco: a concordance score for chromosome conformation capture experiments using random walks on contact map graphs. Bioinformatics 34, 2701–2707 (2018).

28. Yan, K.-K., Yardimci, G. G., Yan, C., Noble, W. S. & Gerstein, M. Hic-spector: a matrix library for spectral and reproducibility analysis of hi-c contact maps. Bioinformatics 33, 2199–2201 (2017).

29. Sauria, M. E. & Taylor, J. Quasar: quality assessment of spatial arrangement reproducibility in hi-c data. BioRxiv 204438 (2017).

30. Bartkuhn, M. & Renkawitz, R. Long range chromatin interactions involved in gene regulation. Biochimica et Biophys. Acta (BBA)-Molecular Cell Res. 1783, 2161–2166 (2008).

31. Peng, J. et al. An end-to-end heterogeneous graph representation learning-based framework for drug-target interaction prediction. Briefings bioinformatics (2021).

32. Zhao, T., Hu, Y., Valsdottir, L. R., Zang, T. & Peng, J. Identifying drug–target interactions based on graph convolutional network and deep neural network. Briefings bioinformatics (2020).

33. Maćkiewicz, A. & Ratajczak, W. Principal components analysis (pca). Comput. & Geosci. 19, 303–342 (1993).

34. Zhang, R., Zou, Y. & Ma, J. Hyper-sagnn: a self-attention based graph neural network for hypergraphs. arXiv preprint arXiv:1911.02613 (2019).

35. Kipf, T. N. & Welling, M. Semi-supervised classification with graph convolutional networks. arXiv preprint arXiv:1609.02907 (2016).

36. Chen, J., Ma, T. & Xiao, C. Fastgcn: fast learning with graph convolutional networks via importance sampling. arXiv preprint arXiv:1801.10247 (2018).

37. Zitnik, M., Agrawal, M. & Leskovec, J. Modeling polypharmacy side effects with graph convolutional networks. Bioinformatics 34, i457–i466 (2018).

38. Wang, Y., Yao, H. & Zhao, S. Auto-encoder based dimensionality reduction. Neurocomputing 184, 232–242 (2016).

39. Van Trees, H. L. & Bell, K. L. Improved bounds on the local meansquare error and the bias of parameter estimators. (2007).

40. Nair, V. & Hinton, G. E. Rectified linear units improve restricted boltzmann machines. In Icml (2010).

41. Karlik, B. & Olgac, A. V. Performance analysis of various activation functions in generalized mlp architectures of neural networks. Int. J. Artif. Intell. Expert. Syst. 1, 111–122 (2011).

42. Kingma, D. P. & Ba, J. Adam: A method for stochastic optimization. arXiv preprint arXiv:1412.6980 (2014).

43. Rumelhart, D. E., Hinton, G. E. & Williams, R. J. Learning representations by back-propagating errors. nature 323, 533–536 (1986).

44. Arthur, D. & Vassilvitskii, S. k-means++: The advantages of careful seeding. Tech. Rep., Stanford (2006).

45. Ramani, V. et al. Massively multiplex single-cell hi-c. Nat. methods 14, 263–266 (2017).

46. Flyamer, I. M. et al. Single-nucleus hi-c reveals unique chromatin reorganization at oocyte-to-zygote transition. Nature 544, 110–114 (2017).

47. Hubert, L. & Arabie, P. Comparing partitions. J. classification 2, 193–218 (1985).

48. Vinh, N. X., Epps, J. & Bailey, J. Information theoretic measures for clusterings comparison: Variants, properties, normalization and correction for chance. The J. Mach. Learn. Res. 11, 2837–2854 (2010).

49. Rosenberg, A. & Hirschberg, J. V-measure: A conditional entropy-based external cluster evaluation measure. In Proceedings of the 2007 joint conference on empirical methods in natural language processing and computational natural language learning (EMNLP-CoNLL), 410–420 (2007).

50. Fowlkes, E. B. & Mallows, C. L. A method for comparing two hierarchical clusterings. J. Am. statistical association 78, 553–569 (1983).

51. Oja, E. & Yuan, Z. The fastica algorithm revisited: Convergence analysis. IEEE transactions on Neural Networks 17, 1370–1381 (2006).

52. Nagano, T. et al. Cell-cycle dynamics of chromosomal organization at single-cell resolution. Nature 547, 61–67 (2017).

53. Lee, D. D. & Seung, H. S. Learning the parts of objects by non-negative matrix factorization. Nature 401, 788–791 (1999).

54. Wall, M. E., Rechtsteiner, A. & Rocha, L. M. Singular value decomposition and principal component analysis. In A practical approach to microarray data analysis, 91–109 (Springer, 2003).

