## Supplementary Document for "A novel framework for single-cell Hi-C clustering based on graph-convolution-based imputation and two-phase-based feature extraction"

### 1 Introduction to compared methods

**Introduction of scHiCluster.** This method uses linear convolution to replace each element in the contact matrix with the weighted average value of itself and its surrounding elements. Then it uses random walk with restart algorithm to capture the information in the contact map. In order to reduce the deviation caused by uneven sequence coverage, this method only retains the top 20% interactions after the imputation steps. Finally, the processed contact matrix is projected to the shared low-dimensional space. The dimensionality reduction method used in this method is PCA.

**Introduction of Decay.** Decay is the contact decay curve method. The original contact matrix of each cell is converted to $G \in \mathbb{R}^{n \times n}$ by $log_2$. The feature vector $G_d^{'} \in \mathbb{R}^{1 \times n^2}$ of each cell is calculated by equation (1), which represents the contact ratio of each distance. Then the characteristics of $q$ different cells are concatenated to matrix $G^{'} \in \mathbb{R}^{q \times n}$, and finally, PCA is performed to extract the feature representation of the cells.

$$G_d^{'} = \frac{\sum_j G_{j,j+d}}{\sum_{i,j} G_{i,j}} \tag{1}$$

**Introduction of PCA、NMF、FastICA and SVD.** PCA, NMF, FastICA and SVD are four types of dimension reduction methods. We take PCA as an example to introduce how to generate the embedding for cells. Firstly, the method transforms the original contact matrix $G$ of each cell by $log_2$ and then reshapes it to $G_v \in \mathbb{R}^{1 \times n^2}$. The new matrices $G^{'} = \mathbb{R}^{q \times n}$ are concatenated by matrices from all cells, $q$ is also the number of cells. Then, PCA reduces the dimension of the contact matrix for each chromosome and concatenates the chromosome feature vector. Finally, the method applies another PCA to extract the final feature representation of all chromosomes for a cell. For NMF, FastICA and SVD, we take a similar process to generate the embedding of cells.

### 2 Supplementary figures

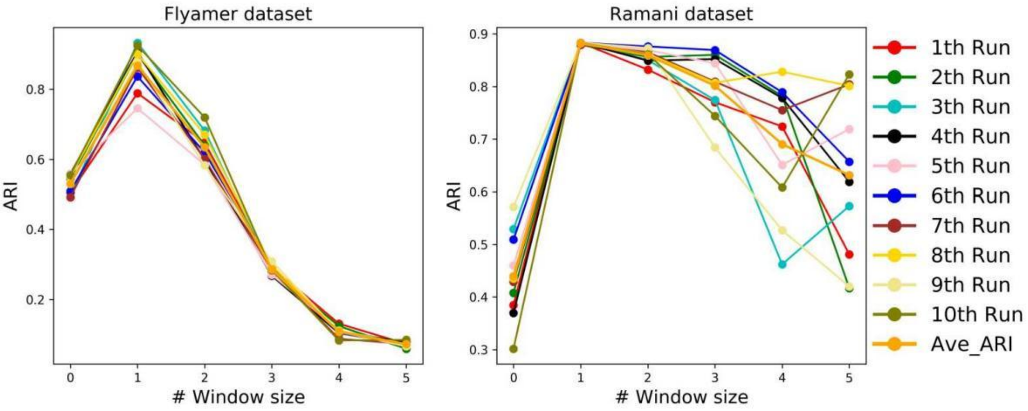


**Figure S1.** The impact of the parameter of window size w on ARI. In genomic neighbor-based imputation step, we use linear convolution to impute contact matrix. In this step, the parameter of window size is w, the filter F size is m×m (where m=2w+1). In our work, window size w is set to 1. Figure S1 shows the effect of window size on ARI.

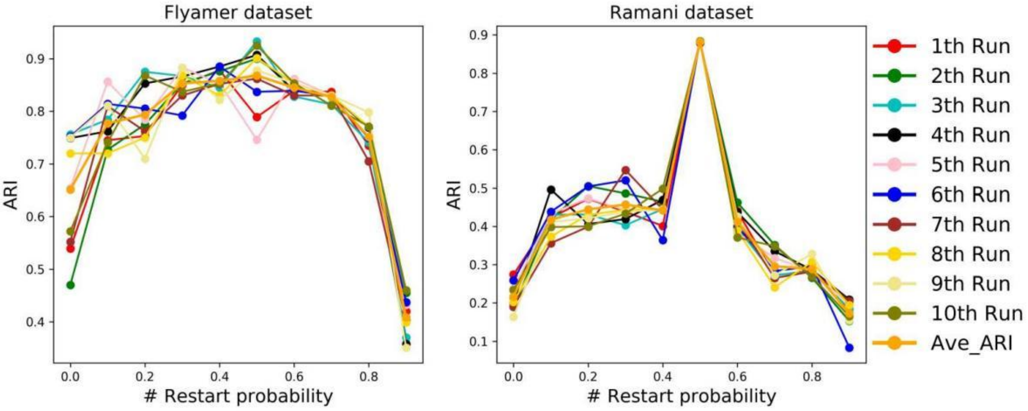


**Figure S2.** The impact of restart probability on ARI. In random walk with restart-based imputation step, we use random walk with restart to impute the contact matrix. In this step, we discuss the effect of restart probability. Figure S3 shows the effect of restart probability size on ARI. In our work, the parameter restart probability p is set to 0.5.

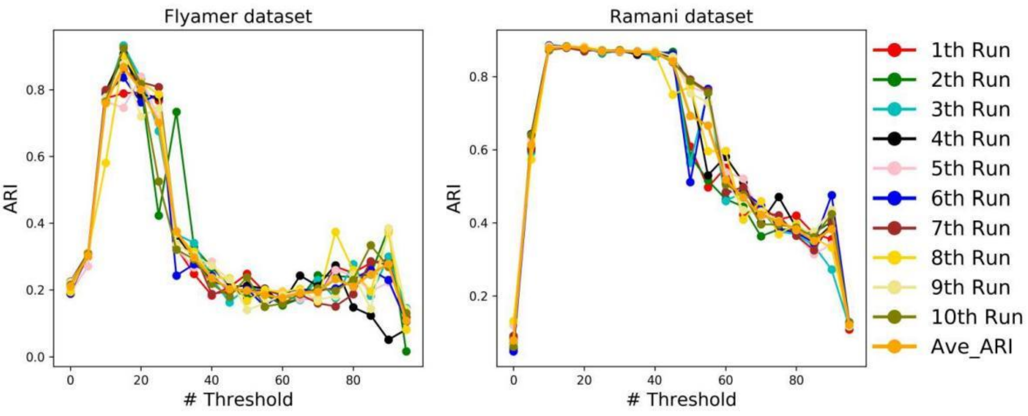


**Figure S3.** The impact of threshold t on ARI. In the step of selecting the top interaction, we convert the matrix to a binary matrix by threshold t. In our work, the threshold t is set to 15, which means that we only select elements in the matrix that are greater than the 85th percentile. Figure S3 shows the effect of the size of t on ARI.

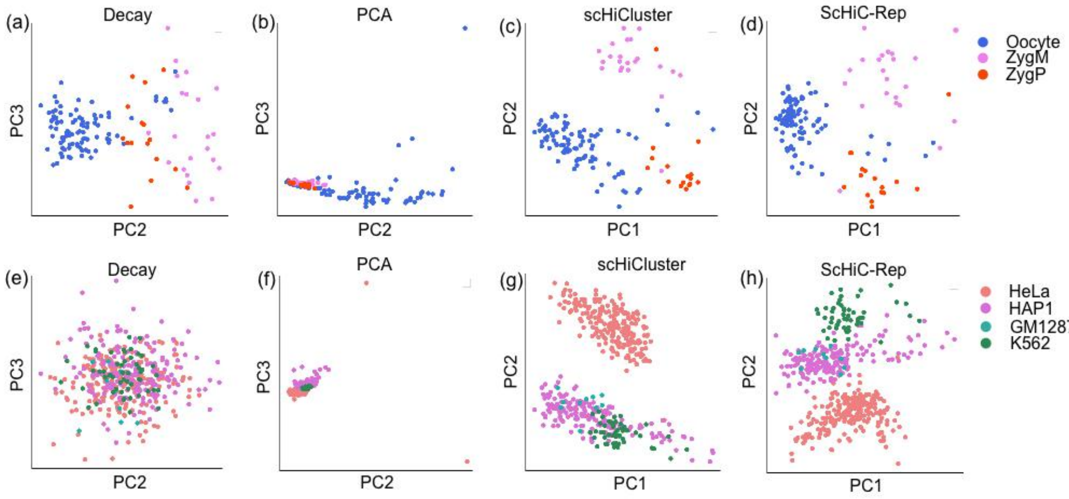


**Figure S4.** Visualization of the learned embedding based on Decay, PCA, scHiCluster and ScHiC-Rep by PCA. (a)-(d) are visualization for Flyamer dataset. (a): Projection of embedding vector for Decay into PC2 and PC3 results in separation of Oocyte from the remaining two cell lines, but weak separation of ZygM and ZygP. (b): The visualization of the three cell lines based on PCA is very compact, but there are no obvious boundaries among the three cell lines. (c): Three cell lines are separated relatively well in the visualization based on scHiCluster, but the boundary between Oocyte cell line and ZygP cell line is not obvious. (d): Three cell lines are separated relatively well in the visualization based on ScHiC-Rep, and ScHiC-Rep makes Oocyte cell line more compact compared with scHiCluster. (e)-(h) are visualization for Ramani dataset. (e): A circular pattern in the visualization based on Decay, where the single-cell of the four cell lines are mixed together. (f): The visualization of the four cell lines based on PCA is very compact except for two HAP1 cells, but there are also no obvious boundaries among the four cell lines. (g): Projection of embedding vector for scHiCluster into PC1 and PC2 results in separation of HeLa from the remaining three cell lines, but weak separation of HAP1, GM12878 and K562. (h): Four cell lines are separated relatively well in the visualization based on ScHiC-Rep. ScHiC-Rep separates K562 cell line from HAP1 cell line and GM12878 cell line compared with scHiCluster.

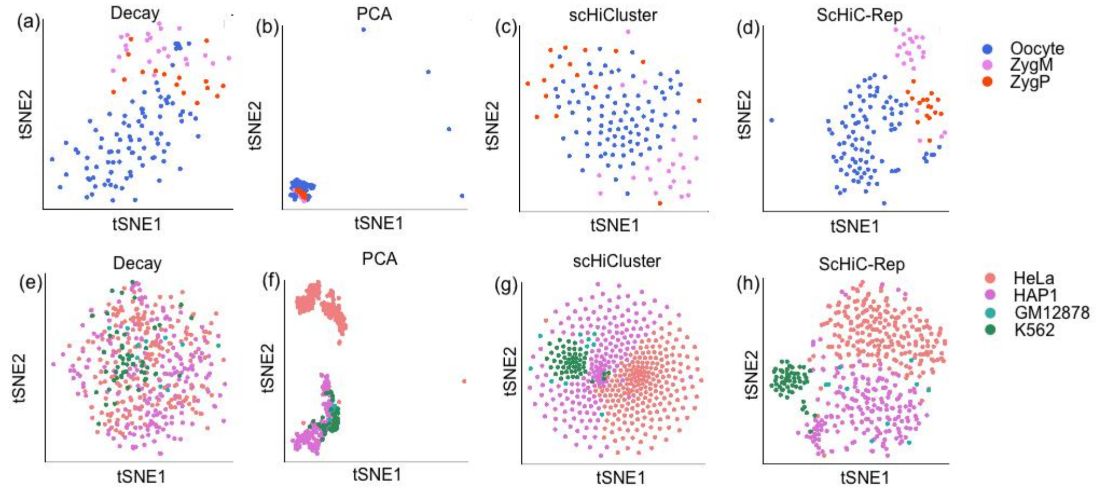


**Figure S5.** Visualization of the learned embedding based on Decay, PCA, scHiCluster and ScHiC-Rep by tSNE. (a)-(d) are visualization for Flyamer dataset. (a): The visualization for Decay does not have boundaries among the three cell lines. (b): The visualization of the three cell lines based on PCA is very compact, but there are no obvious boundaries among the three cell lines. (c): The visualization based on scHiCluster does not have boundaries among three cell lines. (d): Three cell lines are separated relatively well in the visualization based on ScHiC-Rep, and ScHiC-Rep makes three cell lines more compact compared with scHiCluster. (e)-(h) are visualization for Ramani dataset. (e): A circular pattern in the visualization based on Decay, where the single-cells of the four cell lines are mixed. (f): The visualization of the four cell lines based on PCA is very compact, but there are also no obvious boundaries among the three cell lines: HAP1 cell line, GM12878 cell line and K562 cell line. (g): The visualization for scHiCluster only presents these four cell lines in a circular pattern, and there is only a clear boundary among these four cell lines. (h): Four cell lines are separated relatively well in the visualization based on ScHiC-Rep. ScHiC-Rep also separates K562 cell line from HAP1 cell line and GM12878 cell line compared with scHiCluster.

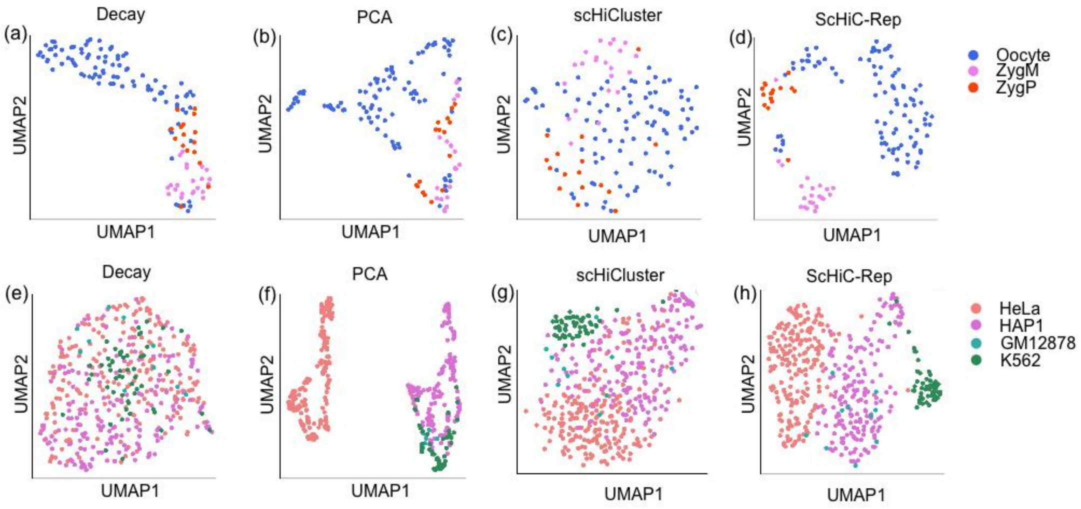


**Figure S6.** Visualization of the learned embedding based on Decay, PCA, scHiCluster and ScHiC-Rep by UMAP. (a)-(d) are visualization for Flyamer dataset. (a): The single-cell for three cell lines are clustered together separately in the visualization for Decay, but it does not have boundaries among the three cell lines. (b): The visualization of the three cell lines based on PCA has no obvious boundaries among the three cell lines. (c): The visualization based on scHiCluster does not have boundaries among three cell lines. (d): Three cell lines are separated relatively well in the visualization based on ScHiC-Rep, and there are three apparent boundaries. (e)-(h) are visualization for Ramani dataset. (e): A circular pattern in the visualization based on Decay, where the single-cell of the four cell lines are mixed. (f): The visualization of the four cell lines based on PCA is very compact, but there are also no obvious boundaries among the three cell lines: HAP1 cell line, GM12878 cell line and K562 cell line. (g): The single-cell for four cell lines are clustered together separately in the visualization for scHiCluster, and there is no clear boundary among these four cell lines. (h): Four cell lines are separated relatively well in the visualization based on ScHiC-Rep. Projection of embedding vector for ScHiC-Rep into UMAP1 and UMAP2 results in separation of HeLa and K562 from the remaining two cell lines, but weak separation of HAP1 and GM12878.

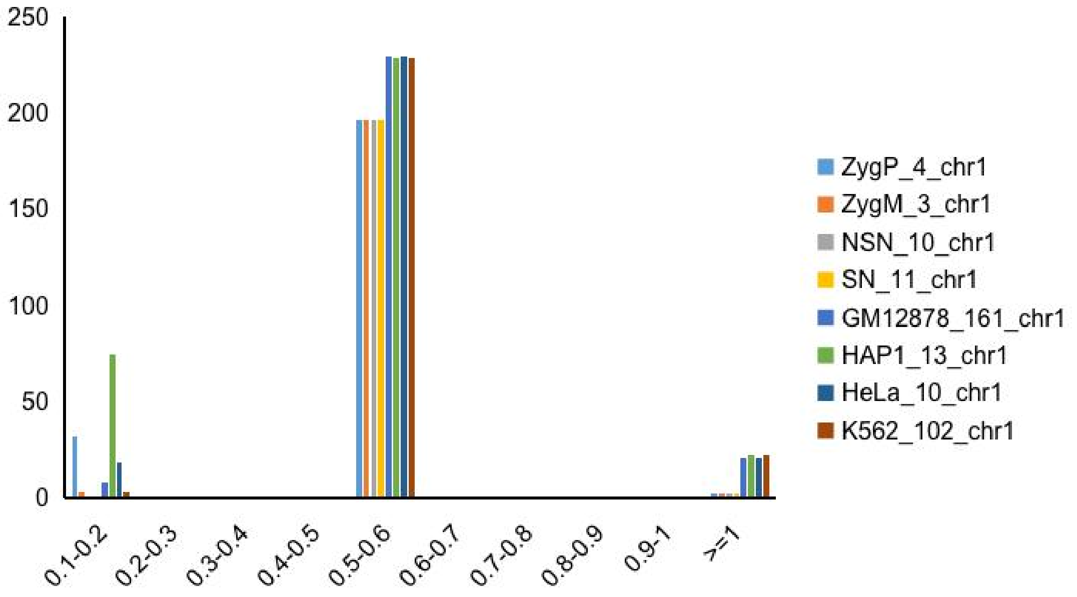


**Figure S7.** Visualization of the distribution for the number of contacts of the original contact matrix of chromosome 1 for each cell line is compared with the distribution for the number of contacts after random walk with restart-based imputation. Cells are randomly selected from Flyamer dataset and Ramani dataset. Since the size of elements in the matrix mainly gather at [0, 0.1], so we only visualize 10 intervals of (0.1, 0.2], (0.2, 0.3], etc. The specific distribution of data can be found in Table S8.

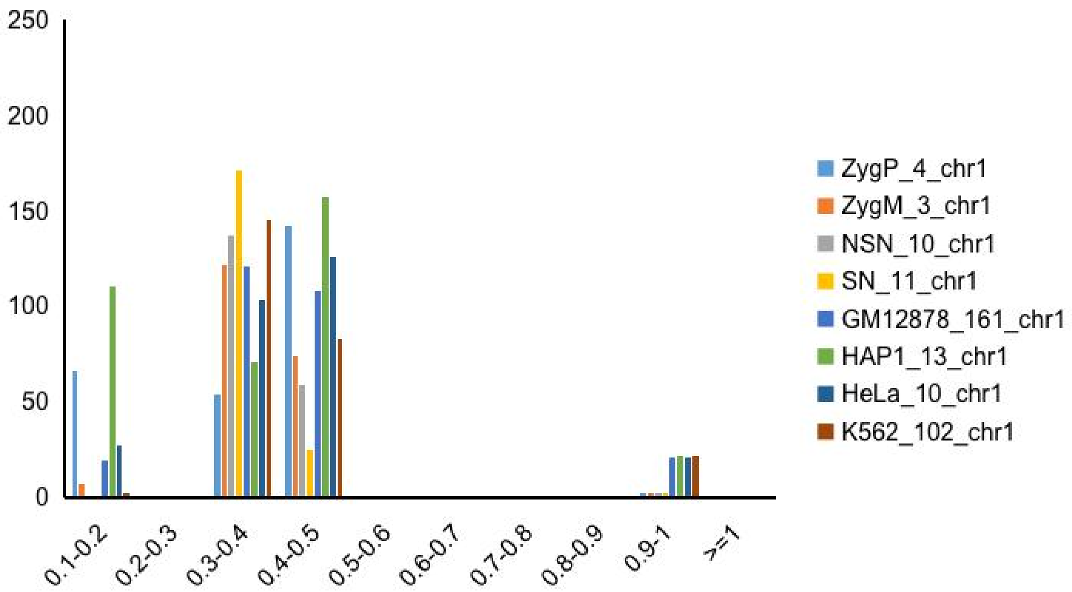


**Figure S8.** Visualization of the distribution for the number of contacts of the original contact matrix of chromosome 1 for each cell line is compared with the distribution of the number of contacts after graph convolution-based imputation. Cells are randomly selected from Flyamer dataset and Ramani dataset. The figure above includes 10 intervals of (0.1, 0.2], (0.2, 0.3], etc. A new interval has been added to the distribution for the data compared with Figure S7, which may be the information which ignores in random walk with restart-based imputation step. The specific distribution for data can be found in Table S9.

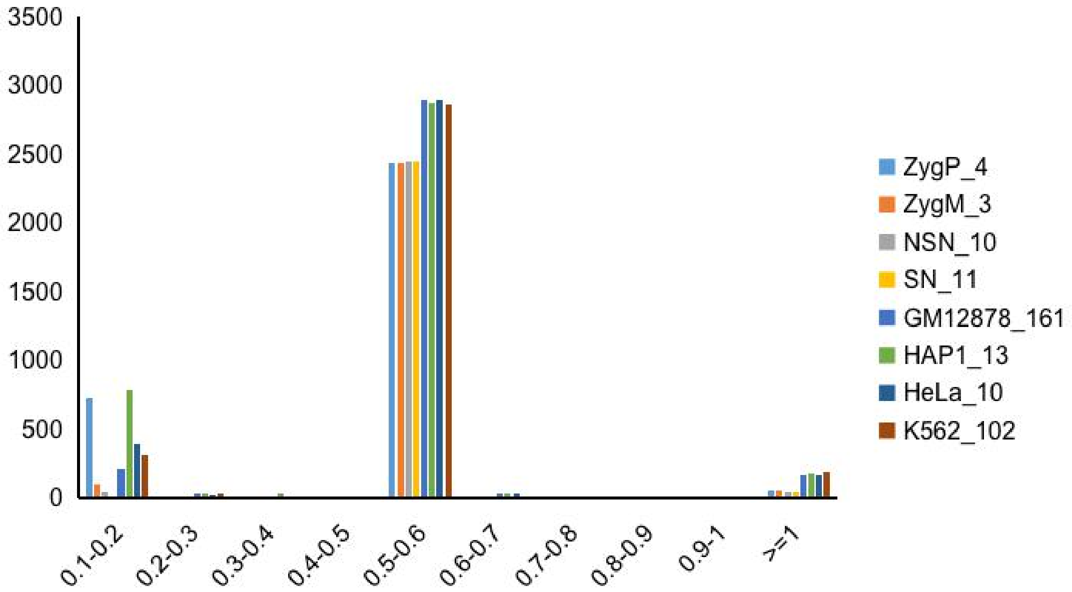


**Figure S9.** Visualization of the distribution for the number of contacts of the original concatenated contact matrix of a cell for each cell line is compared with the distribution for the number of contacts after random walk with restart-based imputation. Cells are randomly selected from Flyamer dataset and Ramani dataset. Since the size of elements in the matrix mainly gather at [0, 0.1], so we only visualize 10 intervals of (0.1, 0.2], (0.2, 0.3], etc. The specific distribution for data can be found in Table S10.

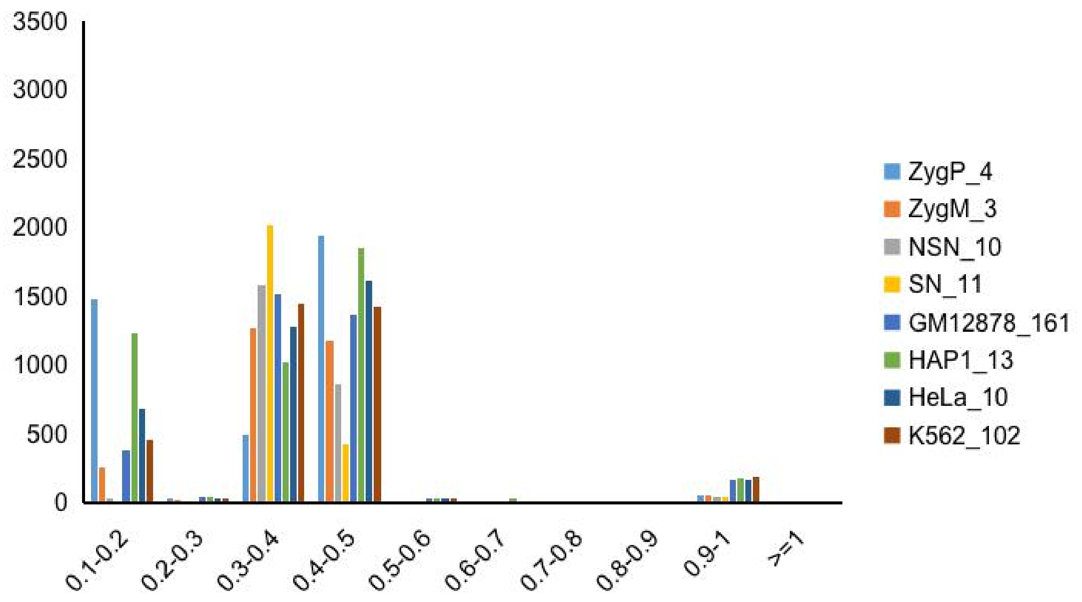


**Figure S10.** Visualization of the distribution for the number of contacts of the original concatenated contact matrix of a cell for each cell line is compared with the distribution for the number of contacts after graph convolution-based imputation. Cells are randomly selected from Flyamer dataset and Ramani dataset. The figure above includes 10 intervals of (0.1, 0.2], (0.2, 0.3], etc. A new interval has been added to the distribution of the data compared with Figure S9, which may be the information which ignores in random walk with restart-based imputation step. The specific distribution of data can be found in Table S11.

### 3 Supplementary table

**Table S1.** The number of cells from Ramani dataset and Flyamer dataset after two steps of filtering. We perform quality control on the two datasets through two-step filtering. In the first step, the contact number of input cells in this model should be greater than 5000 to avoid too few contact numbers of cells. In the second step, assuming that the chromosome length of the input cell in this model is x Mbp, the number of contacts should be greater than x, and all chromosomes of the cell should meet this condition to avoid too few contacts of chromosomes. The number of cell contacts is counted as non-diagonal interacting pairs in the intrachromosomal matrix. Table S1 shows the number of cells from Flyamer dataset and Ramani dataset after two steps of filtering. NSN and SN are two types of Oocytes representing non-surrounded nucleolus and surrounded nucleolus, respectively. ZygP and ZygM are paternal and maternal alleles of zygotes, respectively. Flter1 refers to the criterion that there should be more than 5,000 contacts per cell. Filter2 refers to the criterion that the number of contacts on a chromosome of length x Mbp should be greater than x.

| Dataset | #Cell type | #Cells in total | #Cells after Filter1 | #Cells after Filter2 |
|---|---|---|---|---|
| Ramani | HAP1 | 254 | 229 | 186 |
| | HeLa | 269 | 262 | 217 |
| | GM12878 | 582 | 22 | 10 |
| | K562 | 326 | 72 | 54 |
| | Total | 1431 | 585 | 467 |
| Flyamer | NSN | 40 | 30 | 30 |
| | SN | 76 | 64 | 64 |
| | ZygP | 24 | 18 | 16 |
| | ZygM | 31 | 23 | 22 |
| | Total | 171 | 135 | 132 |

**Table S2.** The optimal hyperparameters of autoencoder for Flyamer dataset calibrated using a grid search procedure. The hyperparameter adjustment process involves learning rate LR, the size of epoch E, learning rate decay β, the number of units in the first hidden layer $H_1$, the number of units in the second hidden layer $H_2$ and the number of units in the output layer O. We choose LR ∈ {0.0001, 0.00001, 0.00002, 0.00003, 0.00004, 0.00005, 0.00006, 0.00007, 0.00008, 0.00009}, E ∈ {200, 300, 400, 500, 600}, β ∈ {0, 0.001, 0.0001}, $H_1$ ∈ {250, 220, 200, 180, 160, 140, 120}, $H_2$ ∈ {120, 110, 100, 90, 80, 70, 60, 50} and O ∈ {40, 32, 24, 16} on the development set of these hyperparameters the best setting.

| Hyperparameter | Selected values |
|---|---|
| LR | 0.0001 |
| E | 500 |
| β | 0 |
| $H_1$ | 220 |
| $H_2$ | 50 |
| O | 16 |

**Table S3.** The optimal hyperparameters of autoencoder for Ramani dataset calibrated using a grid search procedure. The hyperparameter adjustment process involves learning rate LR, the size of epoch E, learning rate decay β, the number of units in the first hidden layer $H_1$, the number of units in the second hidden layer $H_2$ and the number of neurons in the output layer O. We choose LR ∈ {0.0001, 0.00001, 0.00002, 0.00003, 0.00004, 0.00005, 0.00006, 0.00007, 0.00008, 0.00009}, E ∈ {500, 550, 600, 650, 700, 750, 800}, β ∈ {0, 0.001, 0.0001, 0.00001}, $H_1$ ∈ {1000, 900, 800}, $H_2$ ∈ {600, 500, 400, 300} and O ∈ {100, 80, 60, 40, 30, 20} on the development set of these hyperparameters the best setting.

| Hyperparameter | Selected values |
|---|---|
| LR | 0.00004 |
| E | 800 |
| β | 0.00001 |
| $H_1$ | 900 |
| $H_2$ | 500 |
| O | 100 |

**Table S4.** The ARI, NMI, HM and FM of seven methods on Flyamer Dataset. Bolded numbers are the best performance in each category.

| | ARI | NMI | HM | FM |
|---|---|---|---|---|
| FastICA | 0.0270 | 0.0353 | 0.0169 | 0.7292 |
| NMF | 0.0711 | 0.1026 | 0.1172 | 0.4875 |
| PCA | 0.1980 | 0.2156 | 0.2377 | 0.5820 |
| SVD | 0.3661 | 0.4635 | 0.5356 | 0.6566 |
| scHiCluster | 0.6536 | 0.6322 | 0.6925 | 0.8268 |
| Decay | 0.7049 | 0.5644 | 0.5970 | 0.8572 |
| ScHiC-Rep | **0.8049** | **0.7166** | **0.7520** | **0.9058** |

**Table S5.** The ARI, NMI, HM and FM of seven methods on Ramani Dataset. Bolded numbers are the best performance in each category.

| | ARI | NMI | HM | FM |
|---|---|---|---|---|
| FastICA | 0.0327 | 0.0391 | 0.0445 | 0.3447 |
| NMF | 0.1221 | 0.0889 | 0.0866 | 0.4961 |
| PCA | 0.7377 | 0.7404 | 0.6218 | 0.8588 |
| SVD | 0.7461 | 0.6757 | 0.7671 | 0.7104 |
| scHiCluster | 0.8713 | 0.8433 | 0.8776 | 0.9199 |
| Decay | 0.3015 | 0.2918 | 0.3363 | 0.6213 |
| ScHiC-Rep | **0.8822** | **0.8583** | **0.8913** | **0.9264** |

**Table S6.** The sparsity for the original contact matrix of chromosome 1 for each cell line is compared with the sparsity after three steps of imputation. Cells are randomly selected from Flyamer dataset and Ramani dataset (The number behind the cell line represents the ID of the cell. For example, ZygP_4_chr1 represents chromosome 1 of the 4-th cell of the ZygP cell line). Though graph convolution-based imputation does not reduce the sparsity of contact matrix, this step changes the data distribution for contact matrix. The changes in Table S8 and Table S9.

| #sparsity | #original | #after Genomic neighbor-based imputation | #after Random walk with restart-based imputation | #after Graph convolution-based imputation |
|---|---|---|---|---|
| ZygP_4_chr1 | 0.9869 | 0.9040 | 0.0200 | 0.0200 |
| ZygM_3_chr1 | 0.9757 | 0.8336 | 0.0200 | 0.0200 |
| NSN_10_chr1 | 0.9549 | 0.7145 | 0.0200 | 0.0200 |
| SN_11_chr1 | 0.9383 | 0.5587 | 0.0200 | 0.0200 |
| GM12878_161_chr1 | 0.9909 | 0.9207 | 0.1606 | 0.1606 |
| HAP1_13_chr1 | 0.9948 | 0.9598 | 0.1679 | 0.1679 |
| HeLa_10_chr1 | 0.9917 | 0.9315 | 0.1606 | 0.1606 |
| K562_102_chr1 | 0.9921 | 0.9213 | 0.1679 | 0.1679 |

**Table S7.** The sparsity for the original concatenated contact matrix of a cell for each cell line is compared with the sparsity after three steps of imputation. Cells are randomly selected from Flyamer dataset and Ramani dataset. Though graph convolution-based imputation does not reduce sparsity for a cell, this step changes the data distribution for contact matrix. The changes in Table S10 and Table S11.

| #sparsity | #original | #after Genomic neighbor-based imputation | #after Random walk with restart-based imputation | #after Graph convolution-based imputation |
|---|---|---|---|---|
| ZygP_4 | 0.9825 | 0.8879 | 0.0364 | 0.0364 |
| ZygM_3 | 0.9704 | 0.8064 | 0.0372 | 0.0372 |
| NSN_10 | 0.9399 | 0.6579 | 0.0327 | 0.0327 |
| SN_11 | 0.9157 | 0.4661 | 0.0320 | 0.0320 |
| GM12878_161 | 0.9854 | 0.8759 | 0.0818 | 0.0818 |
| HAP1_13 | 0.9906 | 0.9267 | 0.0898 | 0.0898 |
| HeLa_10 | 0.9874 | 0.8993 | 0.0804 | 0.0804 |
| K562_102 | 0.9911 | 0.9100 | 0.1444 | 0.1444 |

**Table S8.** The number of contacts for the original contact matrix of chromosome 1 for each cell line is compared with the number of contacts after random walk with restart-based imputation. Cells are randomly selected from Flyamer dataset and Ramani dataset.

| #contacts of chr1 | ZygP_4 | ZygM_3 | NSN_10 | SN_11 | GM12878_161 | HAP1_13 | HeLa_10 | K562_102 |
|---|---|---|---|---|---|---|---|---|
| 0 | 786 | 786 | 786 | 786 | 10038 | 10494 | 10038 | 10494 |
| 0-0.1 | 38188 | 38219 | 38220 | 38220 | 52204 | 51682 | 52194 | 51755 |
| 0.1-0.2 | 32 | 1 | 0 | 0 | 8 | 74 | 18 | 1 |
| 0.2-0.3 | 0 | 0 | 0 | 0 | 0 | 0 | 0 | 0 |
| 0.3-0.4 | 0 | 0 | 0 | 0 | 0 | 0 | 0 | 0 |
| 0.4-0.5 | 0 | 0 | 0 | 0 | 0 | 0 | 0 | 0 |
| 0.5-0.6 | 196 | 196 | 196 | 196 | 229 | 228 | 229 | 228 |
| 0.6-0.7 | 0 | 0 | 0 | 0 | 0 | 0 | 0 | 0 |
| 0.7-0.8 | 0 | 0 | 0 | 0 | 0 | 0 | 0 | 0 |
| 0.8-0.9 | 0 | 0 | 0 | 0 | 0 | 0 | 0 | 0 |
| 0.9-1 | 0 | 0 | 0 | 0 | 0 | 0 | 0 | 0 |
| >=1 | 2 | 2 | 2 | 2 | 21 | 22 | 21 | 22 |

**Table S9.** The number of contacts for the original contact matrix of chromosome 1 for each cell line is compared with the number of contacts after graph convolution-based imputation. Cells are randomly selected from Flyamer dataset and Ramani dataset.

| #contacts of chr1 | ZygP_4 | ZygM_3 | NSN_10 | SN_11 | GM12878_161 | HAP1_13 | HeLa_10 | K562_102 |
|---|---|---|---|---|---|---|---|---|
| 0 | 786 | 786 | 786 | 786 | 10038 | 10494 | 10038 | 10494 |
| 0-0.1 | 38154 | 38213 | 38220 | 38220 | 52193 | 51646 | 52185 | 51754 |
| 0.1-0.2 | 66 | 7 | 0 | 0 | 19 | 110 | 27 | 2 |
| 0.2-0.3 | 0 | 0 | 0 | 0 | 0 | 0 | 0 | 0 |
| 0.3-0.4 | 54 | 122 | 137 | 171 | 121 | 71 | 103 | 145 |
| 0.4-0.5 | 142 | 74 | 59 | 25 | 108 | 157 | 126 | 83 |
| 0.5-0.6 | 0 | 0 | 0 | 0 | 0 | 0 | 0 | 0 |
| 0.6-0.7 | 0 | 0 | 0 | 0 | 0 | 0 | 0 | 0 |
| 0.7-0.8 | 0 | 0 | 0 | 0 | 0 | 0 | 0 | 0 |
| 0.8-0.9 | 0 | 0 | 0 | 0 | 0 | 0 | 0 | 0 |
| 0.9-1 | 2 | 2 | 2 | 2 | 21 | 22 | 21 | 22 |
| >=1 | 0 | 0 | 0 | 0 | 0 | 0 | 0 | 0 |

**Table S10.** The number of contacts for concatenated original contact matrix of a cell for each cell line is compared with the number of contacts after random walk with restart-based imputation. Cells are randomly selected from Flyamer dataset and Ramani dataset.

| #contacts | ZygP_4 | ZygM_3 | NSN_10 | SN_11 | GM12878_161 | HAP1_13 | HeLa_10 | K562_102 |
|---|---|---|---|---|---|---|---|---|
| 0 | 12504 | 12794 | 11248 | 11002 | 39064 | 42836 | 38388 | 68930 |
| 0-0.1 | 328056 | 328395 | 330030 | 330290 | 434943 | 430603 | 435449 | 404984 |
| 0.1-0.2 | 732 | 103 | 14 | 0 | 213 | 783 | 388 | 309 |
| 0.2-0.3 | 0 | 0 | 0 | 0 | 6 | 2 | 1 | 3 |
| 0.3-0.4 | 0 | 0 | 0 | 0 | 0 | 2 | 0 | 0 |
| 0.4-0.5 | 0 | 0 | 0 | 0 | 0 | 0 | 0 | 0 |
| 0.5-0.6 | 2434 | 2433 | 2439 | 2440 | 2890 | 2873 | 2890 | 2861 |
| 0.6-0.7 | 0 | 0 | 0 | 0 | 3 | 5 | 3 | 0 |
| 0.7-0.8 | 0 | 0 | 0 | 0 | 0 | 0 | 0 | 0 |
| 0.8-0.9 | 0 | 0 | 0 | 0 | 0 | 0 | 0 | 0 |
| 0.9-1 | 0 | 0 | 0 | 0 | 0 | 0 | 0 | 0 |
| >=1 | 48 | 49 | 43 | 42 | 160 | 175 | 160 | 192 |

**Table S11.** The number of contacts for concatenated original contact matrix of a cell for each cell line is compared with the number of contacts after graph convolution-based imputation. Cells are randomly selected from Flyamer dataset and Ramani dataset.

| #contacts | ZygP_4 | ZygM_3 | NSN_10 | SN_11 | GM12878_161 | HAP1_13 | HeLa_10 | K562_102 |
|---|---|---|---|---|---|---|---|---|
| 0 | 12504 | 12794 | 11248 | 11002 | 39064 | 42836 | 38388 | 68930 |
| 0-0.1 | 327308 | 328243 | 330011 | 330290 | 434774 | 430137 | 435146 | 404838 |
| 0.1-0.2 | 1474 | 254 | 33 | 0 | 373 | 1233 | 682 | 452 |
| 0.2-0.3 | 6 | 1 | 0 | 0 | 15 | 18 | 10 | 6 |
| 0.3-0.4 | 495 | 1263 | 1577 | 2013 | 1515 | 1016 | 1272 | 1441 |
| 0.4-0.5 | 1939 | 1170 | 862 | 427 | 1371 | 1854 | 1617 | 1418 |
| 0.5-0.6 | 0 | 0 | 0 | 0 | 7 | 8 | 4 | 2 |
| 0.6-0.7 | 0 | 0 | 0 | 0 | 0 | 2 | 0 | 0 |
| 0.7-0.8 | 0 | 0 | 0 | 0 | 0 | 0 | 0 | 0 |
| 0.8-0.9 | 0 | 0 | 0 | 0 | 0 | 0 | 0 | 0 |
| 0.9-1 | 48 | 49 | 43 | 42 | 160 | 175 | 160 | 192 |
| >=1 | 0 | 0 | 0 | 0 | 0 | 0 | 0 | 0 |
